# GAL4 drivers specific for Type Ib and Type Is motor neurons in *Drosophila*

**DOI:** 10.1101/445577

**Authors:** Juan J. Perez-Moreno, Cahir J. O’Kane

## Abstract

The *Drosophila* larval neuromuscular system is extensively used by researchers to study neuronal cell biology, and *Drosophila* glutamatergic motor neurons (MNs) have become a major model system. There are two main Types of glutamatergic MNs, Ib and Is, with different structural and physiological properties at synaptic level at the neuromuscular junction. To generate genetic tools to identify and manipulate MNs of each Type, we screened for *GAL4* driver lines for this purpose. Here we describe *GAL4* drivers specific for examples of neurons within each Type, Ib or Is. These drivers showed high expression levels and were expressed in only few MNs, making them amenable tools for specific studies of both axonal and synapse biology in identified Type I MNs.

## INTRODUCTION

*Drosophila* research has contributed for decades to our understanding of both fundamental neuroscience (Bellen et al., 2010), and neurological disorders (Ozdowski et al., 2015; Tan and Azzam, 2017; Xiong and Yu, 2018). Much fruitfly neuroscience research is performed at the larval neuromuscular junction (NMJ), a well-characterized system with powerful genetic tools and accessible for physiology and cell biology (Menon et al., 2013).

The larval neuromuscular system has a relatively simple pattern that consists, in abdominal hemisegments from A2 to A7, of around 36 motor neurons (MNs) and 30 muscles (Landgraf and Thor, 2006) (**Fig. 1**), with most muscles co-innervated by more than one Type of MN (Hoang and Chiba, 2001; Kim et al., 2009). Depending on the NMJ bouton properties, different Types of MN have been described in *Drosophila* larvae. Type I MNs are excitatory and glutamatergic, and are subdivided into Ib (big) and Is (small). Type II and Type III MNs are neuromodulatory, being respectively octopaminergic and peptidergic. In addition, glutamatergic Type I MNs show different muscle innervation patterns: each Type Ib MN typically innervates one muscle, whereas each Type Is MN typically innervates several muscles (Hoang and Chiba, 2001; Kim et al., 2009). The different Types of Type I MN also differ in their structural and physiological properties at synaptic level (Atwood and Klose, 2009). Type Ib synapses show shorter and less extensive branching, and support tonic (sustained) firing, whereas Type Is synapses show more extensive branching, and higher synaptic vesicle release efficacy per impulse, are more phasic (transient), and a higher proportion of their vesicle pool is readily releasable (Atwood et al., 1997; Atwood and Klose, 2009; Xing and Wu, 2018).

**Fig. 1.**
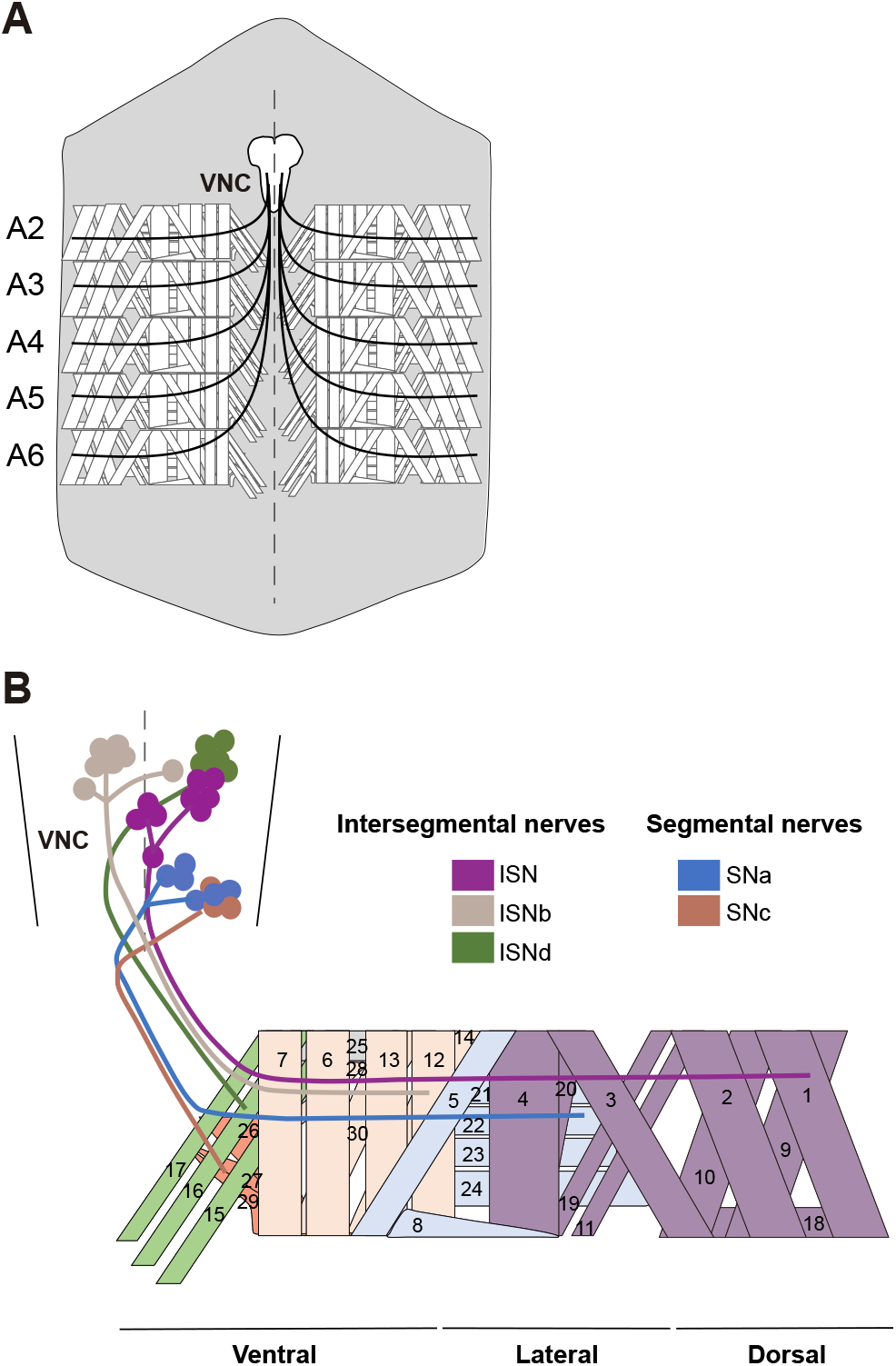
*Drosophila* larval neuromuscular system. **A.** Scheme of a dissected third instar larva showing the neuromuscular system. Only abdominal segments A2-A6 are represented, whose innervation and muscle pattern are identical. The ventral nerve cord (VNC) consists of segmentally repeated neuromeres that are bilaterally symmetrical across the midline (dotted line). Body wall muscles of each hemisegment are innervated by around 30 motor neurons (MNs), whose axons project together from one VNC neuromere, forming a peripheral nerve (black arrow). Anterior is to the top. **B.** Innervation of one of the abdominal hemisegments shown in A. In the VNC, the MN cell bodies project their axons via six main nerve branches: three intersegmental nerves, two segmental nerves, and a transverse nerve (runs along the segment border but has few MNs, so not represented). The muscles innervated by each nerve branch are represented in a lighter version of the color of each branch, and each muscle number is indicated.

To understand the properties of each Type of NMJ synapse, it is important to identify and manipulate different MN terminals independently. A common approach is labelling (typically using anti-Dlg) of the subsynaptic reticulum (SSR), comprising extensive infolding of the postsynaptic cell membrane, and whose amount differs among MN Types (Menon et al., 2013; Zito et al., 1999). However, this approach has several limitations, especially when trying to distinguish different MNs with overlapping branches at the same NMJ: fewer channels available for fluorescence microscopy, especially in live imaging, and potential misidentification of bouton Types in genotypes or environmental conditions that affect SSR or bouton size. An approach to avoid all these limitations would be to use markers based on the genetic identity of the MN.

Using the *GAL4/UAS* system, is possible to express markers or functional proteins specifically in those cells expressing GAL4 (Brand and Perrimon, 1993). While neuromodulatory MNs (Types II and III) are not as extensively studied as excitatory glutamatergic MNs (Ib and Is), specific *GAL4* drivers have been reported for Type II (Stocker et al., 2018) and Type III (Park et al., 2003; Vömel and Wegener, 2007; Koon and Budnick, 2012) MNs. Several useful *GAL4* drivers are expressed in Ib and Is MNs, but in some cases they are also expressed in neuromodulatory MNs (Koon and Budnick, 2012), or in both Type Ib and Is MNs (Fujioka et al., 2003). Other Type I-specific drivers are steroid-activated (Nicholson et al., 2008). In addition, most of the mentioned lines are expressed in multiple MNs, and are therefore less amenable for studies on identifiable axons for which labeling of no more than 2 or 3 MNs would be desirable.

We therefore aimed to identify *GAL4* drivers specific for small numbers of Type Ib or Is MNs. For this, we screened expression patterns in the larval abdominal nerve cord, in some of the neuronal *GAL4* lines generated by the FlyLight project (https://www.janelia.org/project-team/flylight), and identified two glutamatergic GAL4 lines, one specific for a single Type Ib MN, and the other specific for two Type Is MNs. We also identified other potential drivers for neuromodulatory MNs (Type II/III). We propose the two Type I-specific lines as tools of general interest for the *Drosophila* neuroscience community, improving the rigor and the accuracy of the study of both axonal and presynaptic biology.

## RESULTS

### Screening for potential drivers specific for glutamatergic MNs

The FlyLight project has generated around 7,000 transgenic *Drosophila* lines, in each of which expression of *GAL4* is controlled by a different transcriptional enhancer that often drives expression in small subsets of neurons (Pfeiffer et al., 2008; Jennet et al., 2012). To identify drivers specific for different classes of MN, we first reviewed images of the larval central nervous system, for 418 *GAL4* lines listed as driving expression of the *UAS-*m*CD8::GFP* reporter in the abdominal ventral nerve cord (http://flweb.janelia.org/cgi-bin/flew.cgi). We prioritized candidates using several criteria: single or as few as possible cell bodies per neuromere in the VNC; axons visible in the nerves that innervate the body wall musculature (peripheral nerves); moderate or high GFP levels in axons.

We then analyzed selected candidate lines (**Table 1**), using two different reporters to verify the *GAL4* expression levels and distribution: a plasma membrane marker (*UAS-CD4::GFP*) to visualize the whole neuron, including cell body, axonal and presynaptic regions; and an endoplasmic reticulum (ER) marker (*UAS-tdTom::Sec61β*), previously described as continuously distributed throughout the whole neuron (Summerville et al., 2016; Wu et al., 2017). Unless otherwise specified, we refer below to the plasma membrane marker. In addition, we checked the Type of NMJ produced by such cells. Type II presynaptic terminals are smaller and show longer branch length than Type I terminals, while Type III MNs show characteristic elongated or elliptical presynaptic terminals with an intermediate size between Type I and II (Jia et al., 1993). Therefore, we used these properties to choose potential glutamatergic GAL4 lines (**Supp. Fig. S1**), and additionally used anti-Dlg labeling on a subset of lines (**Supp Fig. S2**) to assess the robustness of our identification criteria. We stopped screening once we found a line for each Type I MN (Ib and Is).

**Table 1.**
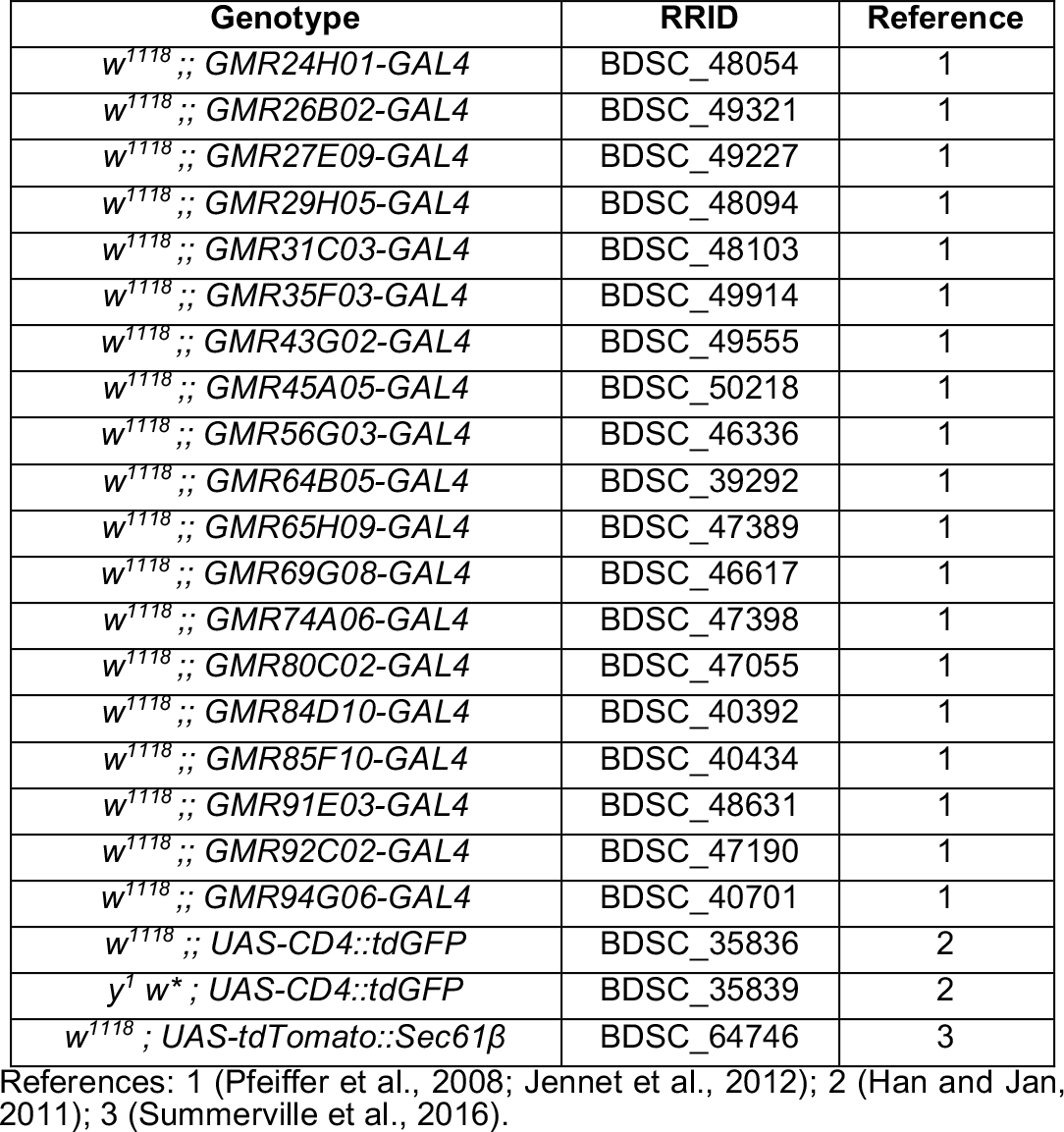
*Drosophila* stocks used in this work.

Using these criteria, we found two lines that appeared to express GAL4 in either Type Is or Type Ib MNs, GMR27E09 and GMR94G06 respectively, described in more detail below. In addition, we found several lines expressing in either Type II or Type III MNs, as well as three lines potentially expressing both in excitatory Type I MNs, and either Type II or Type III modulatory motor neurons (**Supp. Table S1; Supp. Fig. S1, S2**).

### GMR27E09-GAL4 drives expression in two Type Is MNs per hemisegment

*GMR27E09-GAL4* showed expression in two prominent cell bodies per hemineuromere (**Fig. 2 A, B**). These were located close to the midline, and projected their axons towards each peripheral nerve, one ipsilaterally and one contralaterally to the cell body (**Fig. 2 A-B’’**). One of these cell bodies (projecting ipsilaterally) was located in the dorsal region of the VNC, and the other (projecting contralaterally) in the ventral region (**Fig. 2 B-B’’; Supp. Movies S1, S2**). In the peripheral nerve, where both axons run parallel to each other, their paths were too close to distinguish by confocal microscopy in some regions (top panel on **Fig. 2 C**), but we frequently found regions where both axons could be easily distinguished (bottom panel on **Fig. 2 C**). Each axon innervated several internal muscles from a nerve branch found close to the intersegmental region. One of the MNs innervated proximal muscles (ventral and lateral), while the other MN innervated distal muscles (lateral and dorsal) (**Fig. 3 A, B**). Based on these data, we conclude these MNs are respectively part of the ISNb and ISN branches of the intersegmental nerve (Hoang and Chiba, 2001; **Fig.1**).

**Fig. 2.**
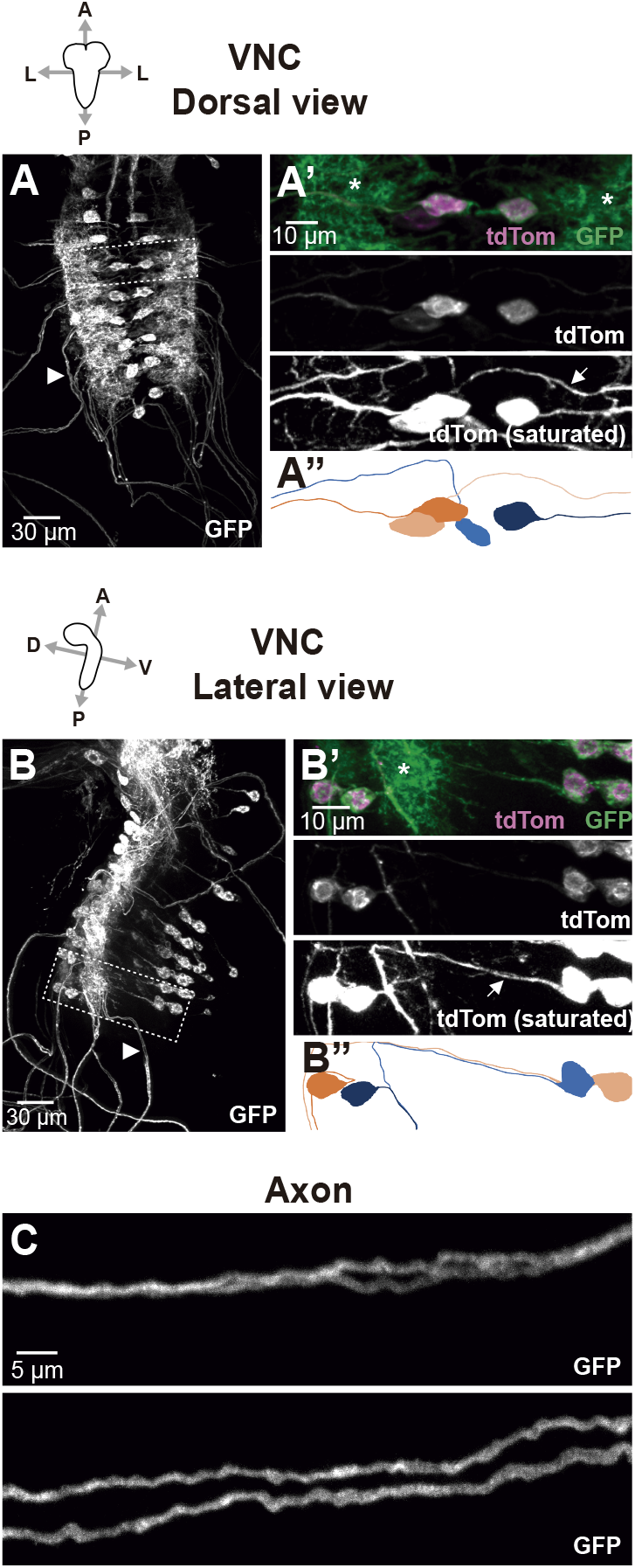
*GMR27E09-GAL4* expression in VNC. Dorsal (**A**) and lateral (**B**) confocal projections showing VNC of larvae expressing *UAS-CD4::GFP* and *UAS-tdTom::Sec61β* under control of *GMR27E09-GAL4.* **A’** and **B’** show magnification of the areas indicated with a dotted line in **A** and **B** respectively. Plasma membrane marker (CD4::GFP) reveals expression of *GMR27E09-GAL4* in two motor neuron cell bodies per hemineuromere (**A’**). Unlike plasma membrane marker, which shows high levels in neuropil (asterisks), axonal projection of each cell body can be easily tracked (arrows) using the ER marker *(tdTom::Sec61β).* The corresponding tracking representations are shown in **A’’** and **B’’**, where neurons with cell bodies in different hemineuromeres are represented in orange and blue. Light and dark versions of the colors are used to distinguish between the two neurons within each hemineuromere. Examples of their axonal projections into the same peripheral nerve are indicated by arrowheads in **A** and **B**. A, anterior; P, posterior; L, lateral; D, dorsal; V, ventral. **C**. In each peripheral nerve, plasma membrane signal reveals regions where both axons mostly overlapped (top panel), or remained apart (bottom panel). Both examples shown in C correspond to different parts of the same peripheral nerve. Anterior and posterior regions are to the left and to the right respectively.

**Fig. 3.**
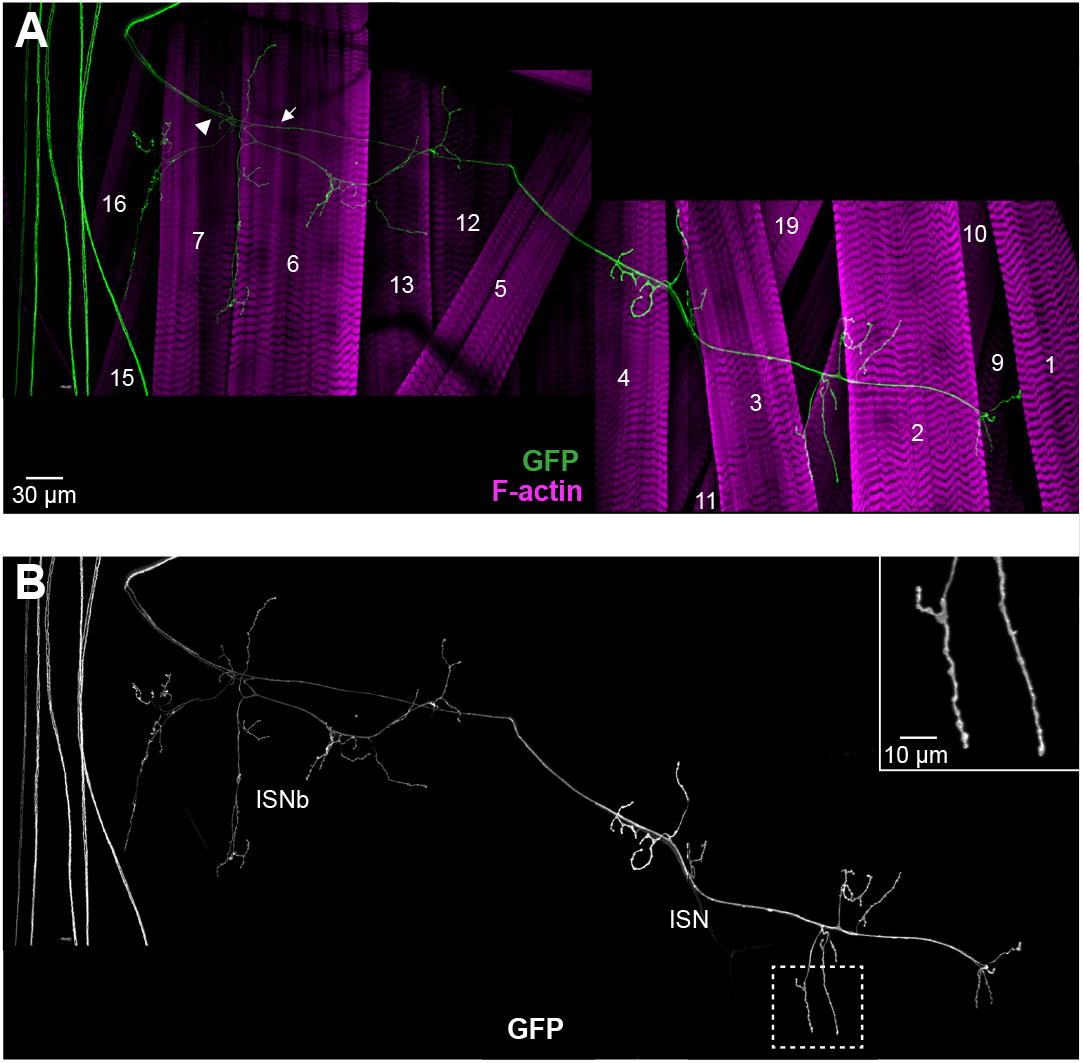
Muscle innervation by MNs expressing *GMR27E09-GAL4*. Composite of several confocal projections showing the NMJ of a whole abdominal hemisegment in a *GMR27E09-GAL4/UAS-CD4::GFP* larva. **A.** Roots of ISNb and ISN nerve branches are indicated by an arrowhead and an arrow respectively. Phalloidin staining of F-actin highlights muscle cells. Recognizable muscles are indicated with the corresponding number. **B.** Single channel image of (**A**) showing GFP expression. Magnification of the area indicated with a broken line is shown in the top right corner. Midline is on the left; anterior is to the top.

The short length of the presynaptic branches and the relatively big size of the presynaptic boutons, suggested that the two MNs expressing *GMR27E09-GAL4* could be Type I or glutamatergic (**Fig. 3 A, B**). To test this possibility, and also distinguish between Type Ib and Is glutamatergic terminals, we double-labeled for *GAL4*-dependent reporter expression and anti-Dlg, whose postsynaptic distribution shows different sizes and levels between Ib and Is boutons (Menon et al., 2014). The NMJ presynaptic terminals of both labeled axons showed detectable levels of Dlg, but did not include Type Ib boutons that showed the biggest and brightest anti-Dlg signals (**Fig. 4 A-C**). Since Dig is absent in Type II and III NMJs (Menon et al. 2013), we conclude that both MNs expressing *GMR27E09-GAL4* are of Type Is.

**Fig. 4.**
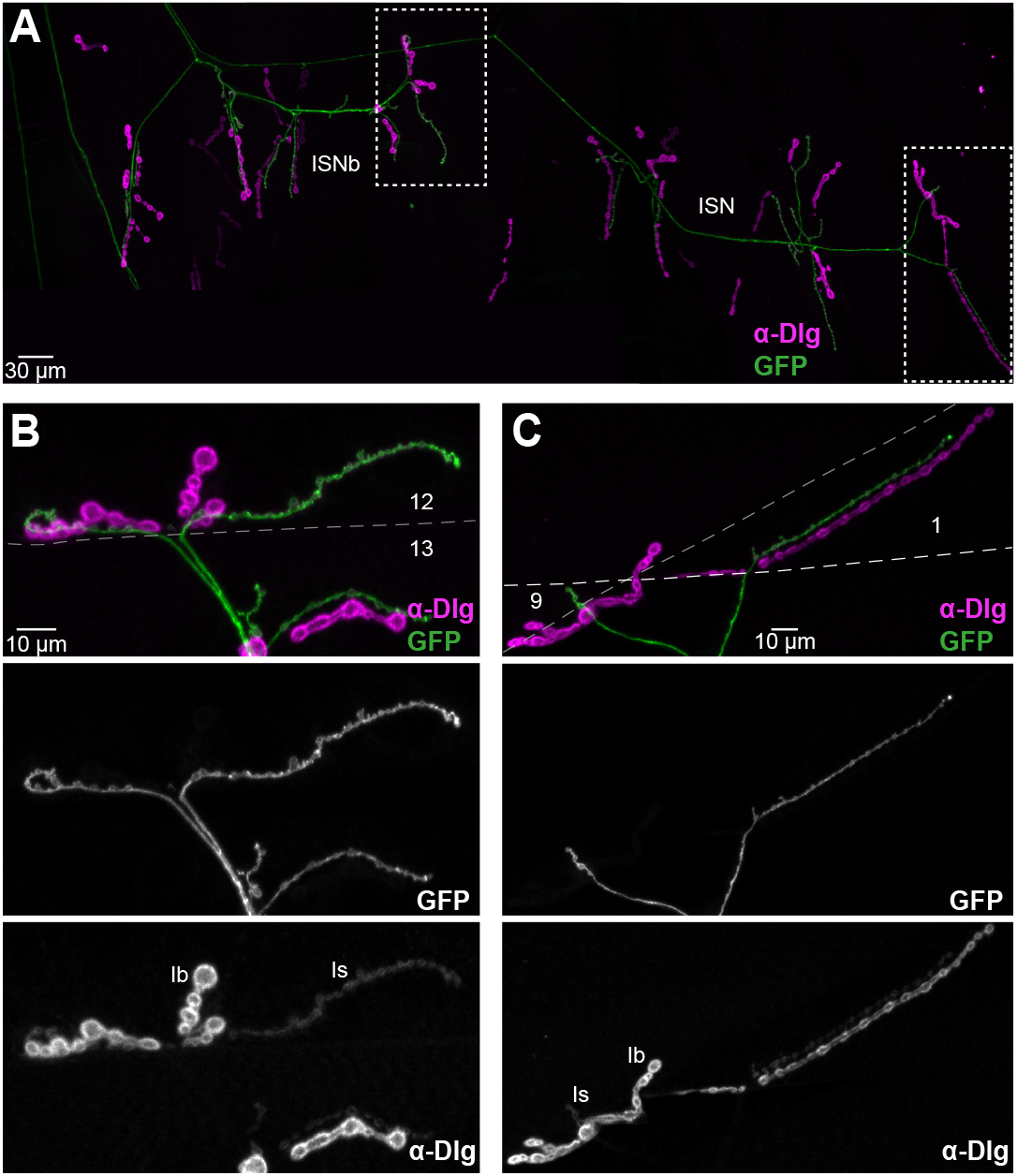
*GMR27E09-GAL4* is expressed in two Type Is MNs. **A.** Composite of several confocal projections showing the NMJ of a whole abdominal hemisegment in a *UAS-CD4::GFP, UAS-tdTom::Sec61β/+; GMR27E09-GAL4/+* larva. Immunostaining of Dlg helps distinguish between Type Ib and Type Is boutons. Midline is on the left; anterior is to the top; positions of major muscles can be inferred by comparison to Fig. 3. Areas inside broken lines are shown at higher magnification in (**B**) and (**C**). Magnified views of NMJs from ISNb nerve on muscles 12/13 (**B**), and ISN nerve on muscles 1/9 (**C**), show that MNs expressing *GMR27E09-GAL4* only produce Is-Type boutons. Dotted lines indicate muscle outlines. In **C**, where both innervated muscles overlapped, edge of muscle 9 is indicated by grey dotted line, while edge of muscle 1 is indicated by white and thick dotted line. Examples of Ib and Is boutons are indicated in the anti-Dlg channel.

At least three Type Is MNs have been described in larval abdominal hemisegments, each innervating multiple muscles from the ISN, SNa and SNb/d branches (Hoang and Chiba, 2001; Kim et al., 2009). Our data suggest that *GMR27E09-GAL4* line is expressed in two of these: ISNb/d-Is, which innervates ventral musculature contralaterally; and ISN-Is (also known as RP2), which innervates lateral and dorsal musculature ipsilaterally (Hoang and Chiba, 2001; Kim et al., 2009).

### GMR94G06-GAL4 drives expression in a single Type Ib MN per hemisegment

Line *GMR94G06-GAL4* was expressed in a single prominent cell body per hemineuromere in the VNC (**Fig. 5 A**). This was located at the dorsal region of the VNC close to the midline, projected its axon ipsilaterally towards the peripheral nerve (**Fig. 5 A’, A’’; Supp. Movies S3, S4**). Therefore, each peripheral nerve contained just a single axon (**Fig. 5 B**). This axon innervated a dorsal muscle (muscle 1) from a nerve branch found close to the intersegmental region (**Fig. 6 A, B**). Therefore, we conclude that the axon follows the ISN branch of the intersegmental nerve (**Fig. 1**). No other innervation of the body wall muscles was detected (**Fig. 6 A, B**).

**Figure 5.**
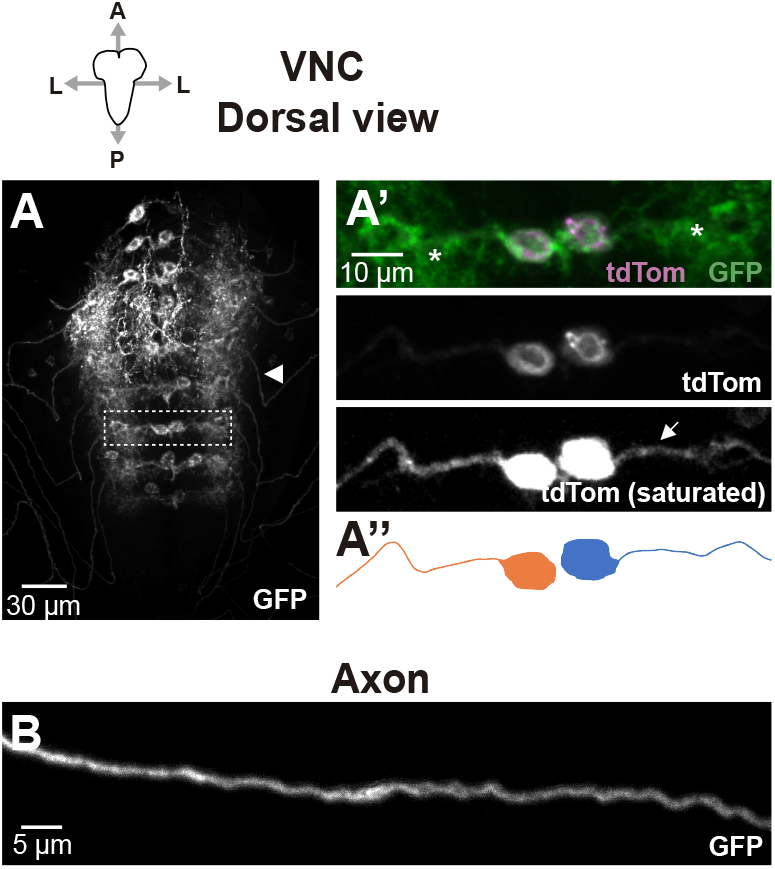
*GMR94G06-GAL4* expression in VNC. **A.** Dorsal confocal projections showing VNC in a *UAS-CD4::GFP, UAS-tdTom::Sec61β/+; GMR94G06-GAL4/+* larva, where plasma membrane marker (CD4::GFP) reveals the expression of *GMR94G06-GAL4* in two cell bodies of muscle- innervating neurons per neuromere. **A’**. Magnification of the areas indicated with a dotted area in **A**. Unlike the GFP plasma membrane marker, which shows high signal levels in the neuropil (asterisks), the axonal projection (arrow) of each cell body can be tracked using ER marker *(tdTom::Sec61β).* The corresponding tracking representation is shown in **A’’**, where neurons innervating different hemisegments are represented in orange and blue. **A**, anterior; P, posterior; L, lateral. **B**. In the peripheral nerve, plasma membrane signal reveals a single axon. Anterior and posterior regions are on the left and the right respectively.

**Figure 6.**
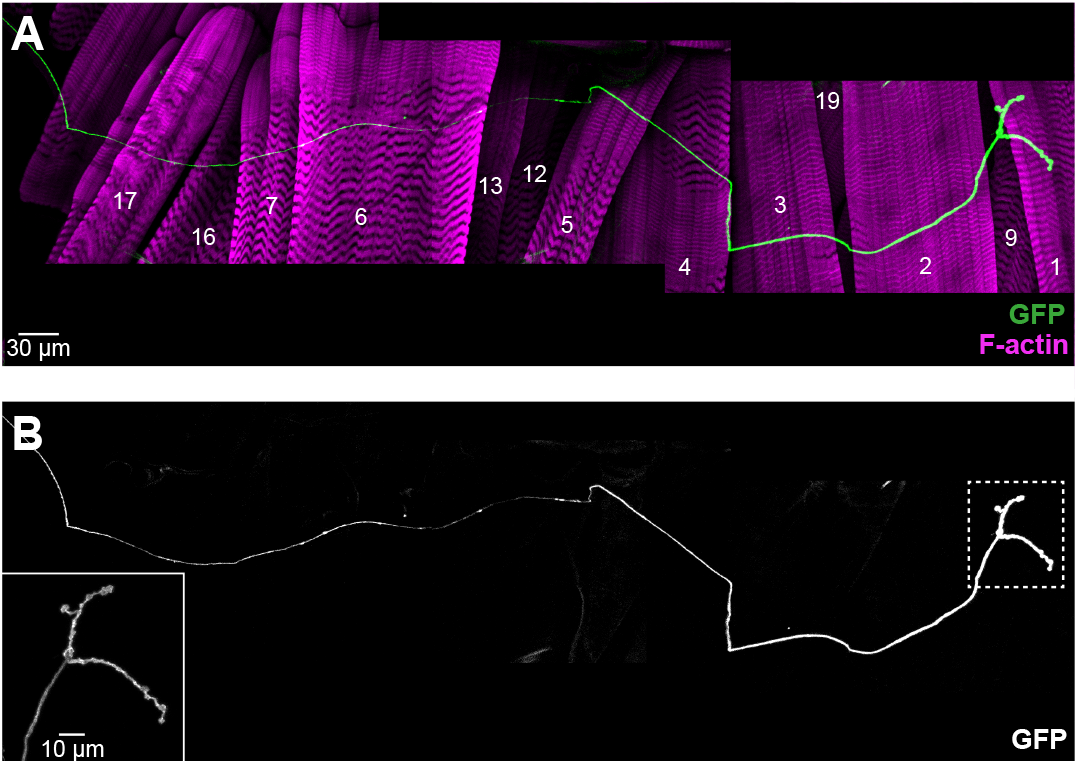
Muscle innervation by MNs expressing *GMR94G06-GAL4*. **A.** Composite of several confocal projections showing the NMJ of a whole abdominal hemisegment in a *GMR94G06-GAL4/UAS-CD4::GFP* larva. Phalloidin staining of F-actin highlights muscle cells. Recognizable muscles are indicated with the corresponding number. **B.** Single channel image of (**A**) showing GFP expression. Magnification of the area indicated with a broken line is shown in the top right corner. Midline is on the left; anterior is to the top.

The large size of the presynaptic boutons, the short length of the NMJ branches, and innervation of a single muscle, suggest that the MN expressing *GMR94G06-GAL4* is a Type Ib glutamatergic neuron. We confirmed this by showing that *GMR94G06-GAL4* drove reporter expression only in presynaptic boutons with high levels of Dlg (**Fig. 7 A, B**). Based on the cell body position, the innervation of muscle 1, and the Type of NMJ (Ib), we conclude that *GMR94G06-GAL4* is specifically expressed in MN1-Ib (also known as aCC) (Hoang and Chiba, 2001; Kim et al., 2009).

**Figure 7.**
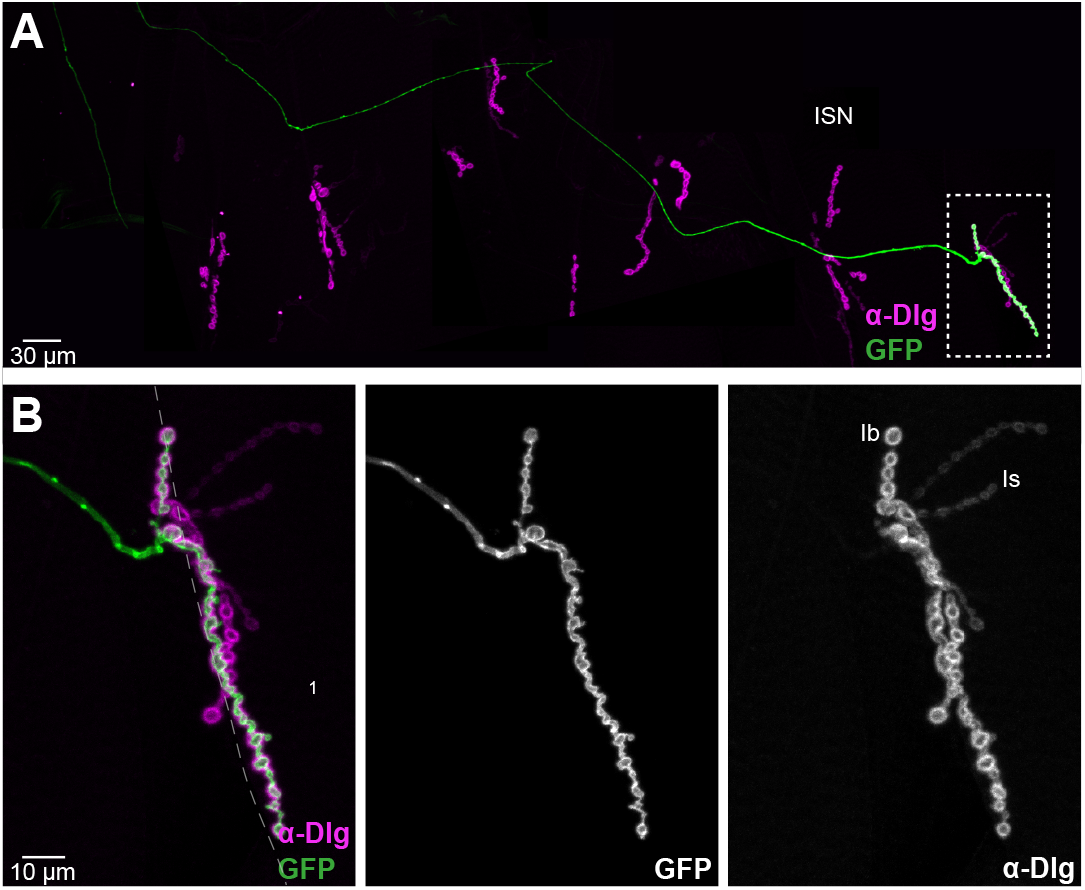
*GMR94G06-GAL4* is expressed in a single Type Ib MN. **A.** Composite of several confocal images showing the NMJ of a whole abdominal hemisegment in a *UAS-CD4::GFP, UAS-tdTom::Sec61β/+; GMR94G06-GAL4/+* larva. Immunostaining against Dlg protein helps distinguish between Type Ib and Type Is boutons. Midline is on the left; anterior is to the top. Area inside broken lines is shown at higher magnification in (**B**). Magnified view of NMJ on muscles 1/9 (**B**) shows that MN expressing *GMR27E09-GAL4* only produces Ib-Type boutons on muscle 1. Dotted lines indicate muscle outlines. Examples of Ib and Is boutons are indicated in the anti-Dlg channel.

### Regulatory regions of GMR27E09-GAL4 and GMR94G06-GAL4 drivers

The MN-specific expression patterns are regulated by enhancers that are used to drive GAL4 expression (Pfeiffer et al., 2008; Jennet et al., 2012). The driver *GMR27E09-GAL4,* expresses GAL4 using a fragment mainly from one of the introns of the *Fmr1* gene, which is present in all recorded transcripts of *Fmr1* (**Fig. 8 A**). In *GMR94G06-GAL4,* the fragment controlling GAL4 expression comes from an intergenic region between two genes of the same family, *dpr4* and *dpr5* (**Fig. 8 B**).

**Figure 8.**
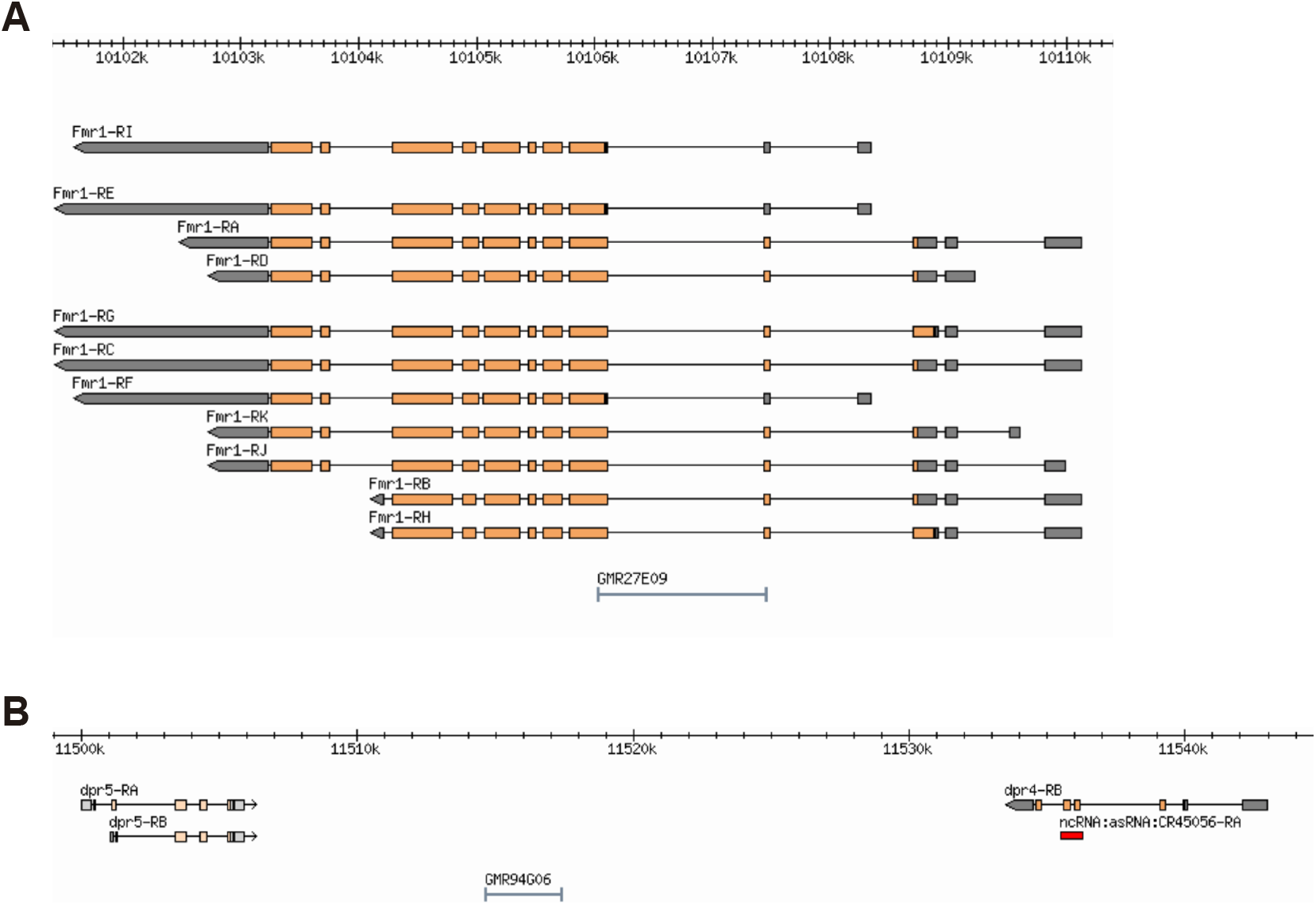
Transcriptional enhancer regions used in *GMR27E09-GAL4* and *GMR94G06-GAL4* constructs. **A, B**. Genomic maps of the regions that contain the fragments that control expression in (**A**) *GMR27E09-GAL4* and (**B**) *GMR94G06-GAL4.* Maps, coordinates, genes and transcripts are from the *Drosophila* Genome Browser (www.flybase.org, version R5.57). Orange regions in transcripts represent coding regions, grey regions represent untranslated regions, and lines represent introns.

## DISCUSSION

### Importance of the identification of GMR27E09-GAL4 and GMR94G06-GAL4 drivers

We have identified and characterized two GAL4 lines specific for different Types of *Drosophila* larval glutamatergic MNs. *GMR27E09-GAL4* is expressed in MN ISNb/d-Is and MN ISN-Is (RP2), while *GMR94G06-GAL4* is expressed in MN1-Ib (aCC). Interestingly, one of the most widely used drivers to analyze specific glutamatergic NMJs is the line *RN2-GAL4* (or *eve-GAL4*), which is expressed in aCC and RP2 (Fujioka et al., 2003; Landgraf et al., 2003). Therefore, the drivers identified here allow to study these same well characterized MNs, aCC and RP2, but separately from each other. The alternative approach of clonal labeling of individual neurons (Roy et al., 2007) requires complex genetics and is not consistent between samples. Here we identified classical GAL4 drivers expressed only in Type Is or Type Ib MNs, which are respectively expressed in two MNs or a single MN per hemisegment, allowing simultaneous axonal and NMJ studies in both fixed or *in vivo* experiments.

### MN identity and regulation of GMR27E09-GAL4 and GMR94G06- GAL4 drivers

The gene expression patterns that govern identity of each MN including its pathfinding and synaptic partners (Landgraf and Thor, 2006), are ultimately regulated by enhancers. The *GMR27E09-GAL4* and *GMR94G06-GAL4* drivers express *GAL4* under the control of genomic regulatory regions from near the *Fmr1*, and *dpr4* or *dpr5* genes respectively.

*Fmr1* encodes an RNA-binding protein, which acts as a neural growth brake regulating RNA trafficking, translation and neuronal excitability, and whose downregulation contributes to Fragile X syndrome in humans (Banerjee et al., 2010). Although *Fmr1* is widely expressed (Wan et al., 2000) and its function is generally required in *Drosophila* larvae MNs (Zhang et al., 2001), the regulatory sequence controlling *GMR27E09-GAL4* (**Fig. 8 A**) drives much more restricted expression than that of *Fmr1.*

The *Dpr* family comprises 21 different genes, which encode neuronal surface receptors required for synapse organization. Several *Dpr* genes are expressed in different subsets of neurons in the *Drosophila* larval VNC, acting as synaptic labels and thus allowing specific synaptic connectivity (Carrillo et al., 2015). Although there is no information available about the expression patterns of the *Dpr4* and *Dpr5* genes located close to the regulatory region in *GMR94G06-GAL4* (**Fig. 8 B**), it is not unexpected that this regulatory region controls expression in a specific MN.

### Future perspectives

*GMR27E09-GAL4* and *GMR94G06-GAL4* are specific drivers for two Type Is MNs and a single Type Ib MN, respectively, per *Drosophila* larvae hemisegment, thus allowing the specific labeling of these Types of MN. This will allow labeling, live imaging, and manipulation of these specific classes of MN, to better understand the biology of the NMJ and its physiologically diverse Types of synapse.

## EXPERIMENTAL PROCEDURES

### Drosophila genetics

All *Drosophila* stocks were obtained from the Bloomington Drosophila Stock Center, and are listed in Table 1.

### Histology and immunomicroscopy

Third instar larvae were dissected in chilled Ca^2+^-free HL3 solution (Stewart et al., 1994), and fixed for 15 min in PBS with 4% formaldehyde. For immunostaining, the dissected preparations were permeabilized in PBS containing 0.1% Triton X-100 (PBT) at room temperature for 1 h. F- actin was stained by incubating dissected samples for 30 minutes at room temperature with Texas Red X-Phalloidin 1:400 (T7471, Thermo Fisher Scientific). For immunostaining, after permeabilization, samples were blocked in PBT with 4% bovine serum albumin for 30 min at room temperature, incubated with primary antibodies overnight at 4°C, and finally incubated with secondary antibodies for 2 h at room temperature. Primary antibody was: mouse anti-Dlg 1:100 (4F3, Developmental Studies Hybridoma Bank; Parnas et al., 2001), and secondary antibody was: goat anti-mouse conjugated to Alexa-647 (A21247, Thermo Fisher Scientific). Processed preparations were mounted in Vectashield (Vector Laboratories), and images were collected using EZ-C1 acquisition software (Nikon) on a Nikon Eclipse C1si confocal microscope (Nikon Instruments, UK). Images were captured using a 40x/1.3NA oil objective.

### Image analysis and Figure preparation

All the microscopy images shown are maximum intensity projections derived from confocal stacks. In the VNC, labeled axonal projections were tracked through sections from cell bodies towards the peripheral nerve, and from the peripheral nerve to the NMJ and muscles. In Fig. 4 and 7, outline of muscles in specific NMJs was identified by using bright- field microscopy. Similarly, the innervation pattern showed in Supp. Table S1. was identified by using bright-field microscopy. All images were opened, analyzed and processed using ImageJ FIJI (https://fiji.sc) (Schindelin et al., 2012). Figures were made using Adobe Illustrator.

## Author contributions

JJPM: Conceptualization, Writing (original draft, reviewing and editing); performed and analyzed all the experiments. CJO’K: Conceptualization, Supervision, Funding acquisition, Project administration, Writing (review and editing).

This article contains supporting information online.

## Acknowledgments

We thank the laboratories of Peter Lawrence and David Glover, and the Developmental Studies Hybridoma Bank, for antibodies, and the Bloomington Drosophila Stock Center for fly stocks.

## SUPPLEMENTARY INFORMATION

**Supplementary Table S1.** GAL4 screen expression summary. Summary of expression data for the *GAL4* drivers screened. Each GAL4 line was crossed with *UAS-Sec61ß::tdTom, UAS-CD4::GFP* except those labeled in grey, which were crossed with *UAS-Sec61ß::tdTom* and immunostained for Dlg. In the Type column “Neuromodulatory” indicates possible Type II and/or Type III MNs.

| Line | MNs | Innervation pattern | Type | Comments |
| --- | --- | --- | --- | --- |
| GMR24H01 | 2 | Ventral muscles | Neuromodulatory | --- |
| GMR26B02 | 2 | Ventral and lateral muscles | Type Is and III | --- |
| GMR29H05 | --- | --- | --- | Homozygous embryonic lethal |
| GMR31C03 | 3 | Ventral, lateral and dorsal muscles | Type II | --- |
| GMR35F03 | 3 | Ventral, lateral and dorsal muscles | Type Ib and II | Much higher expression levels in epidermal cells than in MNs |
| GMR43G02 | 2 | Muscles between ventral and lateral regions | Type III | --- |
| GMR45A05 | 1 | Muscles between ventral and lateral regions | Type III | --- |
| GMR56G03 | 1 | Muscles between ventral and lateral regions | Type III | Low expression levels in the MNs and high in epidermal cells |
| GMR64B05 | 2 | Several ventral and lateral muscles | Type Ib and Neuromodulatory | --- |
| GMR65H09 | 1 | Ventral, lateral and dorsal muscles | Neuromodulatory | MN included in the transversal nerve |
| GMR69G08 | 3 | Ventral, lateral and dorsal muscles | Type II | --- |
| GMR74A06 | 2 | Muscles between ventral and lateral regions | Type III | --- |
| GMR80C02 | 2 | Ventral and lateral muscles | Type II | --- |
| GMR84D10 | 1 | Muscles between ventral and lateral regions | Type III | Low signal levels in axon |
| GMR85F10 | 3 | Several ventral, lateral and dorsal muscles | Type II | --- |
| GMR91E03 | 2 | Lateral muscles | Type II | --- |
| GMR92C02 | 2 | Ventral, lateral and dorsal muscles | Neuromodulatory | --- |

**Supplementary Figure S1.**
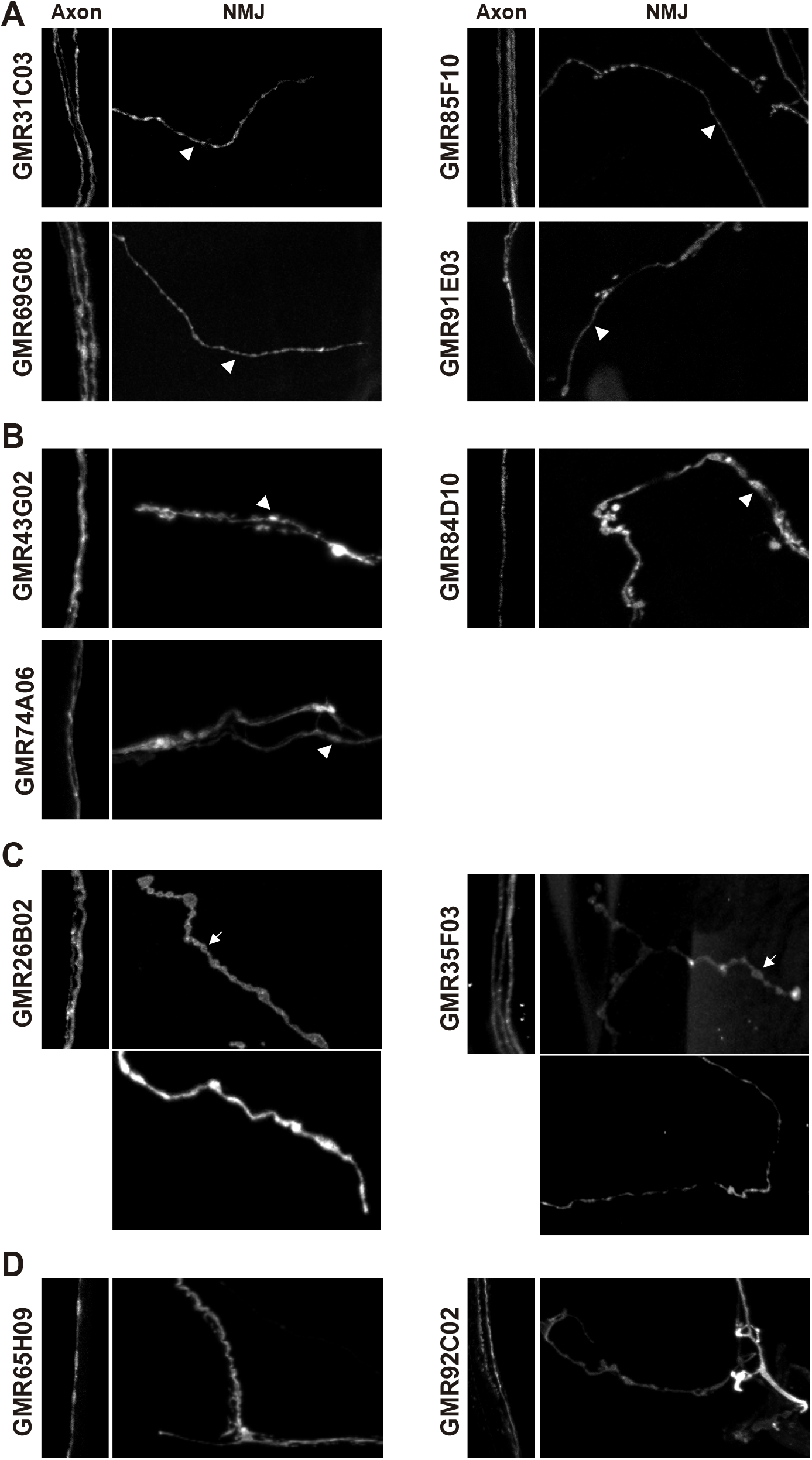
Examples of different MN Types recovered in screen. Confocal projections showing plasma membrane marker (CD4::GFP) in larvae expressing *UAS-CD4::GFP* and *UAS-tdTom::Sec61β* under the control of different GAL4 drivers. Axon images show a region of the peripheral nerve where the MNs expressing the reporter can be distinguished individually, and NMJ images show representative examples of the Types of presynaptic terminal identified. Axon panels: 40×15 μm; NMJ panels: 60×40 μm. **A.** Lines potentially expressed in Type II MNs, which show long NMJ branches and small presynaptic boutons (arrowheads) (Atwood and Klose, 2009). **B.** Lines potentially expressed in Type III MNs, which innervate only a few muscles between the ventral and lateral regions of the hemisegment (Supp. Table S1), and present elliptical-like shaped presynaptic boutons (arrowheads) (Jia et al., 1993; Atwood and Klose, 2009). **C.** Lines potentially expressed in both Type I (top NMJ panels), which show short presynaptic branches with large presynaptic boutons (arrows) (Atwood and Klose, 2009), and neuromodulatory MNs (bottom NMJ panels; Type III and Type II respectively in GMR26B02 and GMR35F03). **D.** Lines that express in an unknown neuromodulatory MN Type (Type II or Type III) but not in Type I MNs.

**Supplementary Figure S2.**
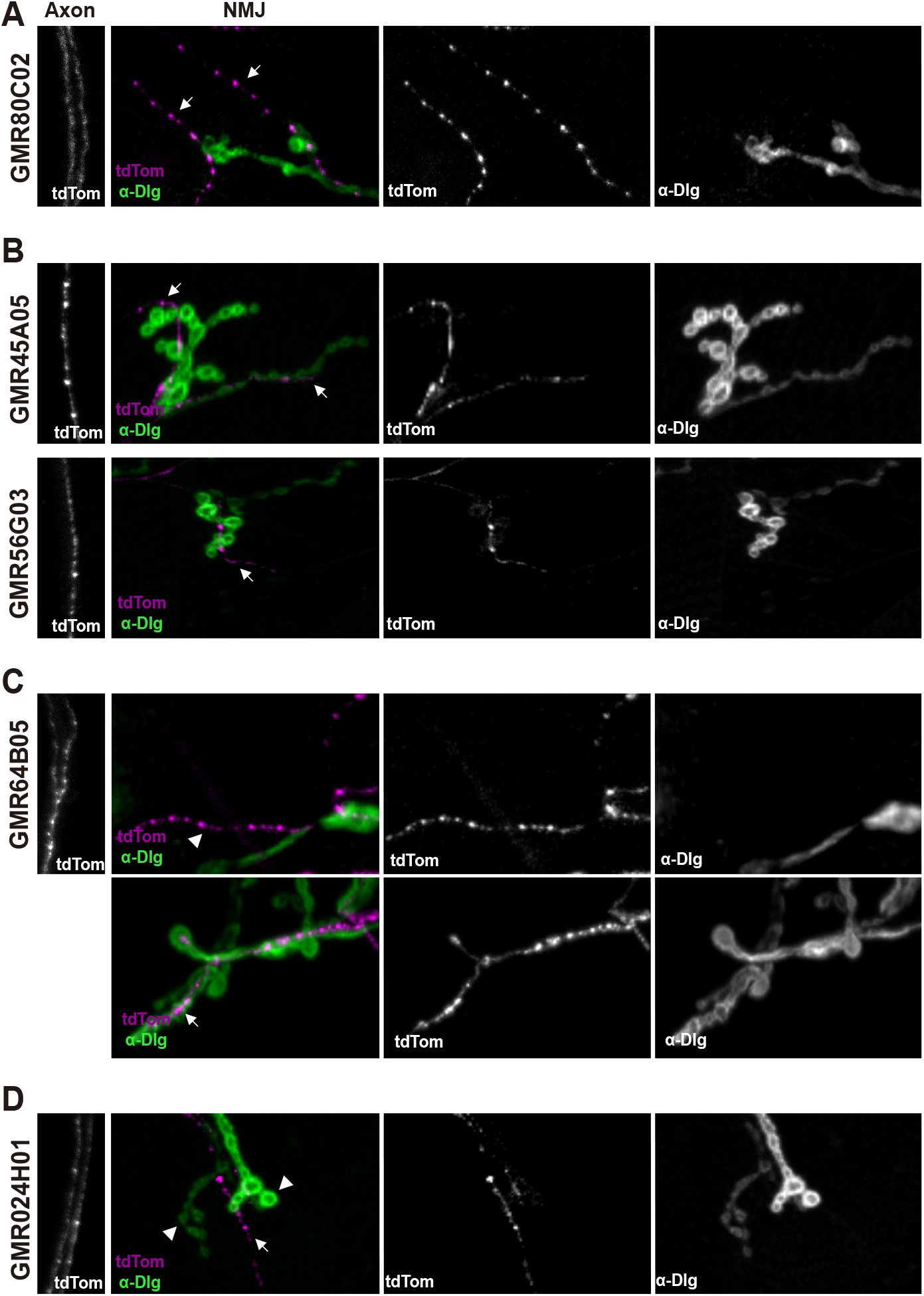
Use of anti-Dlg to confirm MN Types. Confocal projections of larvae expressing *UAS-tdTom::Sec61β* under the control of different GAL4 drivers, organized as in Supplementary Fig. S1. Immunostaining against Dlg protein helps distinguish between different Type of boutons. **A.** Line potentially expressed in Type II MNs, which show long NMJ branches (Atwood and Klose, 2009) and no Dlg signal (arrows) (Menon et al., 2013). **B.** Lines potentially expressed in Type III MNs, which innervate only a few muscles between the ventral and lateral regions of the hemisegment (Supp. Table S1) (Atwood and Klose, 2009), and no Dlg signal (arrows) (Menon et al., 2013). **C.** Line potentially expressed in Type II MNs (top NMJ panel; arrowhead), and in Type Ib (bottom NMJ panel; arrow), which show high Dlg signal (Menon et al., 2013). **D.** Line that expresses in an unknown neuromodulatory MN-Type (arrow) but not in Type I MNs (arrowheads).

**Supplementary Movie S1. VNC showing neuronal plasma membrane labeling by *GMR27E09-GAL4.*** Confocal 3D projection of plasma membrane marker (CD4::GFP) from the VNC of *UAS-CD4::GFP, UAS-tdTom::Sec61β/*+; *GMR27E09-GAL4/*+ larvae. Scale bar, 50 μm.

**Supplementary Movie S2. VNC showing neuronal ER labeling by *GMR27E09-GAL4.*** Confocal 3D projection of ER marker (tdTom::Sec61β) from the same VNC as shown in Supplementary movie S1. Scale bar, 50 μm.

**Supplementary Movie S3. VNC showing neuronal plasma membrane labeling by *GMR94G06-GAL4.*** Confocal 3D projection of plasma membrane marker (CD4::GFP) from the VNC of *UAS-cD4::GFP, UAS-tdTom::Sec61β*/+; *GMR94G06-GAL4/*+ larvae. Scale bar, 50 μm.

**Supplementary Movie S4. VNC showing neuronal ER labeling by *GMR94G06-GAL4.*** Confocal 3D projection of ER marker (tdTom::Sec61β) from the same VNC as shown in Supplementary movie S3. Scale bar, 50 μm.

